# DNA Sequence and Histone Variant H2A.Z Jointly Govern Nucleosome Unwrapping Pathways

**DOI:** 10.64898/2026.06.15.732425

**Authors:** Abhik Ghosh Moulick, Rutika Patel, Tania Rajpersaud, Evgenia N. Nikolova, Sharon M. Loverde

**Affiliations:** Department of Chemistry, College of Staten Island, City University of New York, 2800 Victory Blvd., 6S-238, Staten Island, NY 10314; Ph.D. Program in Biochemistry, Graduate Center, City University of New York, 365 5th Ave, New York, NY 10016, United States; Ph.D. Program in Chemistry, The Graduate Center of the City University of New York, New York, United States; Ph.D. Program in Physics, The Graduate Center of the City University of New York, New York, United States; Institute of Nanotechnology, Karlsruhe Institute of Technology (KIT), Kaiserstraße 12, 76131 Karlsruhe; T.C. Jenkins Department of Biophysics, Johns Hopkins University, Baltimore, MD, 21218

**Keywords:** Coarse-grained model, Nucleosome, SIRAH, All-atom, umbrella sampling, NMR, H2A.Z, nucleosome unwrapping directionality, free energy landscape

## Abstract

Nucleosome unwrapping governs chromatin accessibility and gene regulation, yet the molecular determinants of unwrapping directionality remain poorly understood. Using atomistic and SIRAH coarse-grained umbrella sampling simulations, we show that DNA sequence and histone variant composition jointly tune a directional preference for nucleosome unwrapping. For both the ASP and Widom-601 sequences, unwrapping initiates asymmetrically from a preferred DNA end, with progressive disengagement of the H3 N-terminal tail providing the molecular switch that determines directionality in the Widom-601 system. Substitution of canonical H2A with the variant H2A.Z reverses this directional preference, shifting unwrapping to the opposite DNA end and altering the free energy landscape. SIRAH coarse-grained simulations faithfully reproduce these sequence- and variant-dependent unwrapping pathways and their qualitative free energy features, though quantitative barrier heights differ from atomistic values, identifying a target for further force field refinement. Comparing the H3 tail flexibility from atomistic simulations with published solution NMR amide intensities for two Widom-601 constructs corroborates the fast-timescale tail dynamics and points to a sequence-dependent, microsecond-millisecond exchange component at the H3 tail-core junction. Together, these results establish H3 tail - DNA disengagement, corroborated by NMR data, as a key mechanistic determinant of unwrapping directionality, reveal how a single histone variant substitution can reverse this preference, and validate SIRAH as an efficient framework for large-scale chromatin simulations.

## 1. Introduction

The nucleosome core particle (NCP) is the fundamental building block of chromatin that packages DNA in the cell nucleus. The NCP is composed of approximately 147 base pairs of DNA wrapped around a histone octamer with 1.65 superhelical turns in a left-handed manner. The histone octamer consists of two copies of H3, H4, H2A, and H2B. The NCP, with histone H1 and linker DNA, further assembles into a higher-order, compact chromatin structure^1–6^. The nucleosome is stabilized by the electrostatic interactions between the negatively charged DNA phosphate backbone and the positively charged histone residues. The histone proteins of the histone octamer consist of a globular core and an N-terminal tail. The globular core is an ordered structure composed of three α-helices, and the N-terminal tails are mostly disordered, but they do form secondary structures. The disordered tails are highly charged and flexible with the highest potential for post-translational modifications (PTMs), such as acetylation, phosphorylation, etc due to their higher accessibility to other regulatory proteins^7–13^. The NCP participates in several biological processes, such as transcription, DNA repair, chromosome packaging, and DNA replication^14, 15^.

Epigenetic regulation of chromatin structure and function is influenced not only by histone post-translational modifications (PTMs) but also by histone variants. Canonical histones are incorporated exclusively during the S phase of the cell cycle, where they are added to newly replicated DNA by histone chaperones that collaborate with DNA polymerase. In contrast, histone variants are encoded by genes distinct from those of canonical histones. Their incorporation is not tied to the cell cycle but is instead regulated by specific chaperones or chromatin remodelers that can identify the differences in amino acids between variants and canonical histones. Some variants are highly conserved across species, while others are specific to particular tissues^16, 17^. There are several histone variants that exist for each of the histone proteins, and histone H2A variants exhibit the greatest sequence diversity among all histone variants. The human histone H2A.Z variant is associated with both transcription activation and deactivation^16, 18^.

The NCP undergoes large-scale structural rearrangement during the cell cycles despite being a stable structure. For instance, the NCP disassembly is one of the key steps in DNA replication, and it reassembles after the replication. Small-scale DNA conformational changes, such as partial unwrapping of the nucleosome, occur during chromatin folding and transcription^19–22^. Several experimental studies have shown a dynamic equilibrium between fully wrapped and partially unwrapped DNA, and recent single-molecule long-read footprinting has revealed that more than 85% of nucleosomes in mammalian cells exhibit intranucleosomally accessible DNA, establishing nucleosome distortion as a pervasive in vivo phenomenon rather than a rare exception^23^. Several Förster resonance energy transfer (FRET)^24–27^ and atomic force microscopy (AFM)^28–30^ experimental studies have shown unwrapping and breathing motions of nucleosomal DNA. In addition, several single-molecule force spectroscopy experiments show that the nucleosome unwrapping occurs by starting with the outer turn of the DNA opening at a lower force, followed by the inner turn at a higher force^28, 31^. *Ngo et al.*^29^ showed asymmetric unwrapping of the nucleosome under tension, with one of the DNA tails interacting more strongly with the histone core. Further, circular dichroism (CD)^32, 33^ and NMR^34–36^ studies show that histone tail fluctuations are essential for nucleosome packaging and the formation of higher-order structure. In particular, solution and magic-angle-spinning NMR studies show that the H3 N-terminal tail remains locally disordered within the nucleosome and undergoes exchange between DNA-bound and released conformations, with the strength of this engagement tunable by linker DNA, linker histone binding, and histone post-translational modifications^37–39^. The H3 and H4 N-terminal tails are long and participate in nucleosome packaging, and the removal of these tails increases DNA exposure. Also, the removal of the H2A C-terminal tail increased the unwrapping rate of the nucleosome^36, 40, 41^. The histone H2A.Z variant is known to dynamically decrease nucleosome stability, promoting asymmetric DNA unwrapping and increasing chromatin accessibility. Further, H2A.Z has been shown to facilitate the binding of pioneer transcription factors Sox2 and Oct4 by increasing DNA accessibility and reducing competition with the H3 N-terminal tail, with MD simulations and NMR experiments demonstrating that H2A.Z incorporation leads to more disordered H3 N-terminal tail conformations with fewer DNA contacts, promoting chromatin remodelling^42–49^.

Molecular dynamics (MD) simulation techniques provide atomistic insights into nucleosome dynamics. Several all-atom and coarse-grained simulations have been performed to investigate these dynamics^50–53^. Some studies have specifically focused on sequence-dependent behavior using nucleosomal DNA sequences such as the Widom-601^54^ and the alpha-satellite palindromic (ASP) sequence^55^. Earlier work^56^ from our lab explored distinct unwrapping pathways, where the ASP sequence exhibited loop formation, while the Widom-601 sequence showed large-scale breathing motions based on 12 μs atomistic simulations performed on Anton 2 at a high salt concentration of 2.4 M.

In parallel, coarse-grained (CG) simulation methods have emerged as important tools for studying sequence-dependent nucleosome dynamics^52^. Our lab previously employed the SIRAH CG force field to investigate both sequences, revealing sequence-dependent breathing behavior at the CG level^57^. In addition, de Pablo’s group developed a CG nucleosome model by combining the three-site-per-nucleotide (3SPN) DNA model with the atomic-interaction-based coarse-grained (AICG) protein model, successfully capturing sequence-dependent nucleosome dynamics and exploring repositioning mechanisms via loop propagation or twist diffusion^58^. Takada’s group further used CG modeling to demonstrate two distinct sliding modes depending on the nucleosomal DNA sequence^59^.

Nucleosome wrapping and unwrapping occur on microsecond-to-millisecond timescales, making them challenging to study with conventional all-atom MD simulations as well as CG simulation. Therefore, biasing methods, specifically steered molecular dynamics (SMD) methods and umbrella sampling, have been used to study nucleosome unwrapping. *Takada et al.*^60^ showed the unwrapping process under various salt conditions and found that the unwrapping of nucleosomal DNA is correlated with H3 and H2A tail dynamics. Using the combination of 3SPN DNA and AICG protein coarse-grained model, *Lequieu et al.*^61^ evaluated the tension-dependent free energy landscape and demonstrated that the contributions of histone tails to DNA stability vary between the H3/H4 and H2A/H2B tails. Also, *Zhang et al.*^62^ showed the free-energy surface of nucleosome unwrapping using a coarse-grain model with umbrella sampling and replica exchange, indicating that asymmetric unwrapping of nucleosomal DNA is correlated with histone protein disassembly. *Kono et al.*^63^ performed adaptively biased molecular dynamic (ABMD) simulation to obtain free energy profiles for unwrapping the outer-turn of nucleosomal DNA. Despite these efforts, how DNA sequence and histone variant substitution such as H2A.Z independently and in combination modulate nucleosome unwrapping pathways and free energy landscapes has not been systematically investigated at atomistic level. Furthermore, recent study from our group^57^ employing the SIRAH CG force field have demonstrated its ability to reliably reproduce nucleosome structural dynamics at the base-pair level, showing good agreement with atomistic simulations. In addition, recent studies^64^ have demonstrated the ability of the SIRAH coarse-grained force field to capture specific protein-protein interactions, including those between the histone variant CENP-A nucleosome and its binding partner CENP-N. However, whether SIRAH can faithfully capture sequence and histone variant dependent nucleosome unwrapping pathways and free energy landscapes in direct comparison with atomistic simulations remains to be established.

Here, we perform all-atom and SIRAH coarse-grained steered MD and umbrella sampling simulations across three nucleosome systems, the ASP genomic sequence, the Widom-601 strong positioning sequence, and an H2A.Z variant nucleosome systems, to establish the free energy basis of sequence and variant-dependent unwrapping directionality. We identify progressive H3 N-terminal tail disengagement as the molecular switch governing which DNA end unwraps preferentially in Widom-601, show that H2A.Z substitution reverses this directional preference, and validate SIRAH as a qualitatively reliable framework for exploring these landscapes at reduced computational cost. To link these dynamics to experiment, we compare the per-residue H3 tail fluctuations from our atomistic simulations to recently published NMR amide peak intensities for two nucleosomes carrying Sox2-Oct4 composite sites and either the canonical H2A or the variant H2A.Z histone^49^. The fast-timescale tail response is generally reproduced, whereas a divergence at the flexible hinge^39, 65^ located near the H3 tail-core junction exposes an additional DNA sequence-dependent slow motion. Our atomistic and SIRAH coarse-grained simulations collectively demonstrate that nucleosome unwrapping pathways and their sequence dependence are robustly captured at both levels of resolution. While SIRAH faithfully reproduces the qualitative unwrapping pathways and sequence-dependent behavior observed in atomistic simulations, quantitative differences in free energy barrier heights reflect the inherent limitations of coarse-grained representations. Together, these findings establish SIRAH as a reliable tool for exploring nucleosome unwrapping pathways at reduced computational cost, while highlighting the necessity of atomistic simulations for precise free energy quantification.

## 2. Methods

### 2.1. System preparation

Here, we consider two different sequences of nucleosome core particle (NCP), (i) human l:z-satellite palindromic sequence (ASP) and (ii) the strong positioning ‘Widom-601’ DNA sequence. The initial coordinate for ASP is taken from protein data bank having PDB ID 1KX5^55^. The crystal structure of 1KX5 contain 14 Mn^2+^ ions. Because of absence of force-fields for Mn^2+^, we replaced these ions with Mg^2+^. For Widom-601 sequence, we consider initial coordinate obtained from protein data bank having PDB ID 3LZ0^54^. This crystal structure has missing histone N-terminal tails. So, we modeled these missing tails and other missing residues using Prime of Schrodinger software suite^66, 67^. The H2A.Z nucleosome was modeled using PDB ID: 1F66^68^ by swapping canonical 3LZ0 H2A with H2A.Z from PDB: 1F66^68^ system. The ASP structure mentioned before has been used as the template for this homology modeling purpose. We replaced the 8 Mn^2+^ ions in the crystal structure of homology-modeled 3LZ0 system by Mg^2+^ ion to use available force fields for Mg^2+^ ions.

### 2.2. All atom simulation of the NCP

Both NCP sequences were subjected to all-atom molecular dynamics simulation in the presence 0.15M physiological NaCl salt. The histone proteins were parametrized using the AMBER19ffSB force field^69^ whereas the DNA is parametrized using OL15^70^. The OPC water model^71^ was used as solvent around the NCP in an orthorhombic box. Na^+^ and Cl^-^ ions were parametrized using Joung and Cheetham parameters (2008)^72^ while the Li/Merz compromised parameter set^73^ was used for Mg^2+^ ions. The Lennard-Jones interaction of Na^+^/OPC (OW) was improved according to *Kulkarni et al.* to get better estimation of osmotic pressure^74^.

After parametrization, both systems were minimized using 15000 steps, following both steepest descent and conjugate gradients in the AMBER18 package^75^. Therefore, systems were heated at constant volume, varying the temperature slowly up to 310K. All bonds involving hydrogen atoms were constrained using the SHAKE^76^ algorithm. The heated structures were further equilibrated for 100 nanoseconds (ns) maintaining a constant pressure of 1 Bar using Berendsen barostat^77^ and constant temperature around 310K using Langevin thermostat^77^ with a collision frequency of 1.0 ps. Electrostatics was calculated using the Particle Mesh Ewald (PME)^78^ algorithm with full periodic boundary conditions. A cut-off value of 12□ was used for the van der Waals interaction, while bonded atoms were excluded from the non-bonded atom interactions using a scaled 1-4 value. System specific descriptions are given in **SI Table 1**.

### 2.3. Coarse-grained simulation of the NCP

Next, both NCP systems are simulated using the SIRAH CG forcefield^79^ in the GROMACS 2002.5^80, 81^ package in explicit water. The simulation of the NCPs follows the protocol mentioned in our earlier study^57^. At first, the PDB2PQR server^82^ has been used to set the correct protonation state based on the assumption of neutral pH following the AMBER naming scheme. After mapping into the CG representation, both systems are solvated in a cubic box of SIRAH WT4 water.^83^ The system was neutralized and maintained at 0.15M salt concentration by adding Na^+^ and Cl^-^ ions. The details of the box size, required number of ions, total number of atoms as well as number of solvent atoms are tabulated in **SI Table1**. The systems were energy minimized in two steps, at first protein side chains are energy minimized by keeping restraints on the backbone for 50,000 steps following steepest descent algorithm. Therefore, the whole system was energy minimized for 5000 steps. At the next step, solvent molecules were equilibrated around the complex for 5 ns keeping harmonic restraints on the position of all CG beads. The temperature of the system was maintained at 310K using a V-rescale thermostat. Finally, a 25 ns equilibration was performed to improve the solvation of protein side chains. All simulation parameters were kept the same as in the earlier study of the same system using the SIRAH coarse-grained force field^57^.

### 2.4. Steered MD and umbrella sampling simulation

To construct the free-energy profile of the nucleosome unwrapping we performed steered molecular dynamics (SMD) and umbrella sampling (US)^84–86^ simulations of the NCP using atomistic as well as coarse-grained molecular dynamics simulations. PLUMED^87^ (version 2.8.2) integrated with GROMACS was used to carry out steered molecular dynamics and umbrella sampling simulations. The US method was used following the SMD method using the radius of gyration (R_g_) of DNA as the reaction coordinate^88, 89^, calculated over all DNA atoms. The DNA radius of gyration was selected as the reaction coordinate because it provides a continuous, end-agnostic measure of overall DNA expansion throughout the unwrapping process, and has been employed in prior coarse-grained and atomistic nucleosome umbrella sampling studies^63, 90^, enabling direct comparison with the literature. Critically, because R_g_ does not preferentially bias unwrapping toward either DNA end, the directional preference observed across windows emerges from the underlying sequence and histone composition rather than from the choice of coordinate. The limitation of this approach is that R_g_ alone cannot resolve which end is unwrapping within a given window; directionality is therefore characterized post hoc through H3 tail–DNA contact analysis, SHL0–end distances, and unwrapped base pair counts as described below. Other reaction coordinates used in prior coarse-grained and atomistic studies of nucleosome dynamics, including DNA rotational and translational order parameters^58^, sliding and rotational coordinates^59^, and end-to-end distance, could complement R_g_ in future work to resolve directional unwrapping preferences more directly, building on our recently reported Markov state modeling of histone tail conformational dynamics ^91^.

We consider the equilibrated structure obtained from atomistic simulations to perform further enhanced sampling steps. We performed 2 different simulations i.e. 1KX5 and Widom-601 sequence in physiological salt. System-specific details related to the steered molecular dynamics are tabulated in **SI Table 2**. We extracted representative frames from the SMD trajectory using a spring constant of 10000 kJ mol^-1^ nm^-2^ in such a way as to prepare different US windows, such that the spacing between different windows is 0.05 nm. These configurations were identified as initial structures for the US simulations at each window. Next, we performed molecular dynamics in each window for further 50 ns by implementing a harmonic potential, 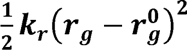 where the force constant ***k_r_*** = 837 kJ mol^-1^ nm^-2^ and 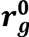 is reference radius gyration of the DNA. We discarded the initial 5 ns of the data for further equilibration. Finally the potential of mean force (PMF) is calculated over the equilibrated trajectory of the US simulation using the weighted histogram analysis method^92^.

In the CG SMD and the US simulations, the equilibrated structures obtained from the CG simulations were used for both sequences. The entire protocol for setting up the SMD simulations and subsequently performing the US simulations using the CG force field was the same as that used in the atomistic simulations. The system-specific details are tabulated in **SI Table 1**. The errors in the PMF plots were estimated by averaging over three different blocks in each window for both the all-atom and CG US simulations.

### 2.5. Unwrapping Features

Features of the unwrapping are analyzed considering the (i) histone-DNA contacts, (ii) dyad (SHL0)-DNA end distance, and (iii) number of unwrapped base pairs using our in-house codes. The contacts are defined by two non-hydrogen atoms from DNA and histone within 4.5 Å following earlier studies^93^. The dyad–DNA distance is defined as the centre-of-mass (COM) distance between either DNA end (terminal 10 bp) and the base pair at the dyad position (see **Figure 1A)**. The unwrapped base pairs are defined as those in which the center of the base pair deviates more than 7 Å from the corresponding base pair in the initial structure.

**Figure 1.**
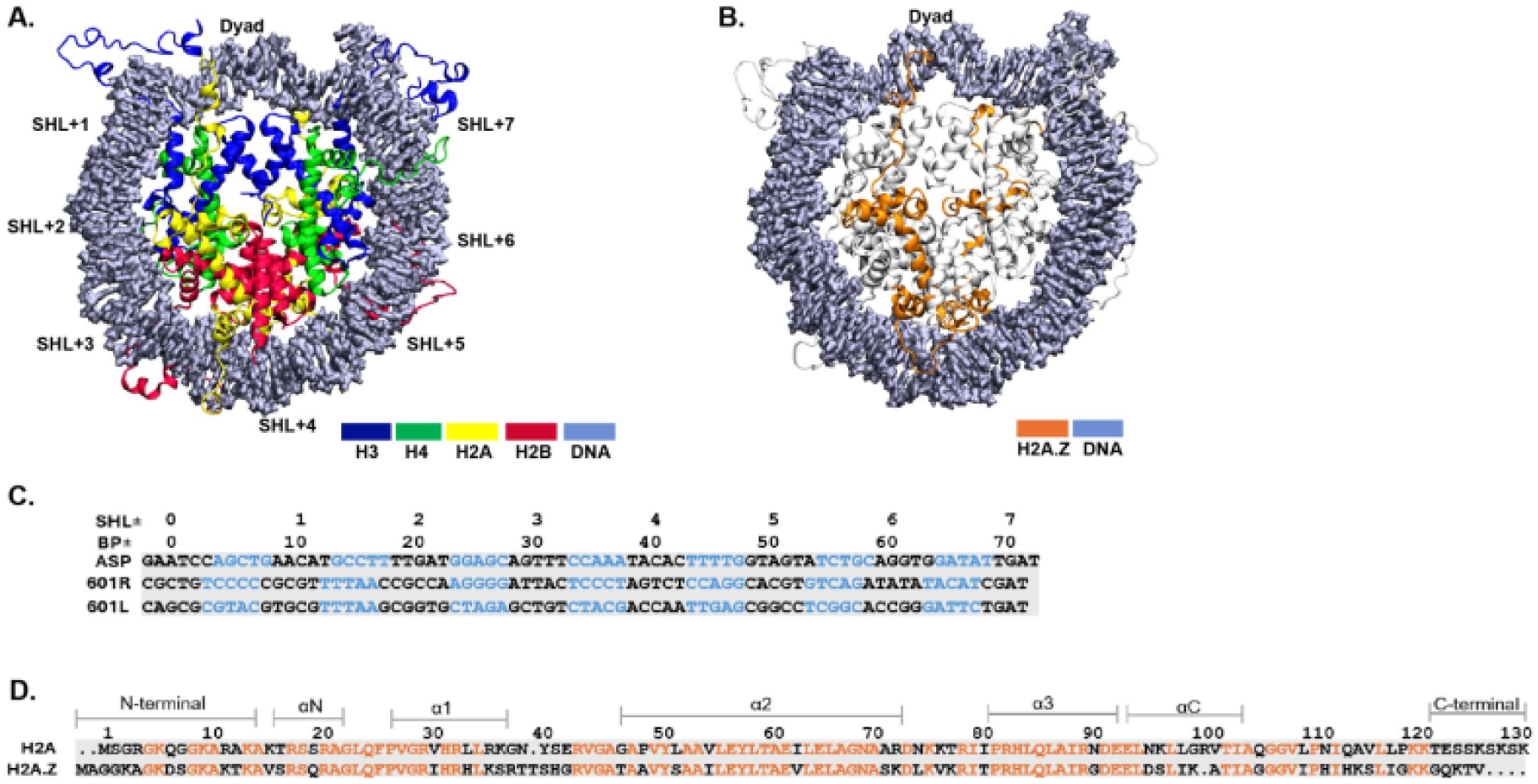
Nucleosome Core Particle (NCP) Structures. **(A)** The crystal structure of NCP (PDB ID: 3LZ0) consists of 145 DNA base pairs wrapped around two copies of the histone proteins H3 (blue), H4 (green), H2A (yellow), and H2B (red). The super helix locations (SHLs) of DNA are labeled on the outside of the DNA. **(B)** The NCP shows the H2A.Z variant, where the H2A.Z (orange) is the only histone swapped as per PDB ID: 1F66, and other histones are the same as the PDB ID: 3LZ0 system. **(C)** The DNA sequences of the two crystal structures used in this study: the human α-satellite sequence (ASP, PDB ID: 1KX5) and the Widom-601 (PDB ID:3LZ0) sequence. For the ASP sequence, only half of the sequence is shown because it is palindromic, whereas for the Widom-601 sequence, both the right (601R) and left (601L) sequences are shown. The blue DNA bases indicate minor grooves, while black DNA bases indicate major grooves. **(D)** The sequence comparison between the human H2A and H2A.Z histone monomers is shown. Identical residues are shown in orange.

### 2.6 Comparison of simulated H3 tail flexibility with NMR amide intensities

Per-residue RMSF of the H3 N-terminal tail (Tail1, residues 1-38) was computed on backbone amide nitrogen atoms for the canonical H2A and H2A.Z Widom-601 (NCP 601) systems from the umbrella-sampling window corresponding to the global PMF minimum. The equilibrated trajectory from this specific window was used to calculate the RMSF values, representing the fluctuations of the H3 N-terminal tail in the most stable conformational state along the unwrapping pathway. The RMSF calculation was performed for backbone C_α_ after removing overall translational and rotational motions by fitting the trajectory to the histone core backbone atoms. The change upon variant incorporation was defined as ΔRMSF = RMSF_H2A.Z_ − RMSF_H2A_. Published H2A.Z/H2A backbone amide peak intensity ratios for the modified Widom NCP 601-62_F_ and NCP 601-62_N_ constructs^49^, containing the mouse Fgf4 (5’ CTTTGTTTGGATGCTAAT) and Nanog (5’ CATTGTAATGCAAAA) Sox2-Oct4 composite motifs on the right side (601-R) spanning approximately SHL+5 to +6.5, were compared with the NCP 601 ΔRMSF values. The strength of association was quantified by the Pearson coefficient (r, linear dependence) and the Spearman rank coefficient (ρ, monotonic dependence), each with two-sided p-values, computed both across all resolved tail residues (n = 28 and 27 for 601-62_F_ and 601-62_N_, respectively) and after excluding residues 31–34 (n = 24 and 23); the latter test isolates the contribution of these residues to the overall trend. Least-squares linear fits were used for visualization, and associations with p < 0.05 were considered significant. Correlation analyses were performed in SciPy^94^ and NumPy^95^ and figures were generated with Matplotlib.

## 3. Results

Here, we perform all-atom and SIRAH coarse-grained steered MD simulations, followed by umbrella sampling (US) for nucleosome systems in the 1KX5^55^ structure that contains a human α-satellite palindromic (ASP) DNA sequence, and the 3LZ0^54^ structure that contains a Widom-601 DNA sequence. The structure of the nucleosome core particle (NCP) consists of DNA wrapped around the histone octamer, as shown in **Figure 1A**. The NCP consists of about 147 DNA base pairs wrapped around two copies of histone proteins H3, H4, H2A, and H2B. The orientation of the DNA base pairs is represented relative to the central DNA base pair, known as superhelical location SHL 0. Each SHL region consists of about 10 base pairs **(Figure 1A)**. The superhelical location is defined where the major groove faces the histone octamer. The first is SHL 0 (at the NCP dyad) and the last is SHL±7. The histone variant H2A.Z has all the histone features similar to the Widom-60l system, except that both H2A histones are replaced with the H2A.Z variant as per PDB ID: 1F66 **(Figure 1B)**. The sequence comparison of the nucleosomal DNA between ASP and Widom-601 is shown in **Figure 1C**. The minor groove at SHL±1.5 in the Widom-601 sequence adopts a narrow conformation, containing a strong positioning TTTAA motif. There is 15% greater GC content in the Widom-601 sequence than in the ASP sequence. For the Widom-601 sequence, the GC content in the right half of the sequence (601-R, TA-poor side) has higher GC content, which makes it more rigid with fewer contacts with the DNA and easier to open it up, as demonstrated by *Ngo et al*^29^. The histone H2A and H2A.Z sequence alignment shows that the C-terminal tail of histone H2A.Z is shorter than that of H2A **(Figure 1D)**.

### 3.1. Nucleosome Unwrapping via Steered MD and Umbrella Sampling Simulation

To explore nucleosome unwrapping, we employed steered MD followed by umbrella sampling, using the DNA radius of gyration as the reaction coordinate. Three sets of all-atom simulations have been carried out for three systems: ASP, Widom-601, and the H2A.Z variant. The systems were first equilibrated and then steered MD simulations were performed. We extracted representative frames from the SMD trajectory to prepare different US windows, spaced 0.05 nm apart. These configurations were identified as the initial structure for US simulation at each window. Next, we performed molecular dynamics in each window for further 50 ns and calculated the potential of mean force (PMF) over the US simulation using the weighted histogram analysis method (WHAM)^92^. The detailed PMFs obtained for all three systems are shown in **Figure 2**. The first minimum in the PMF for the ASP sequence corresponds to the of DNA around 45 Å with further unwrapping, a neighboring minimum appears at 48 Å and 51 Å indicated by vertical dashed lines (**Figure 2A**). The ASP sequenc appears to be unwrapping at the left side of the NCP. Further, the first minimum in the PMF for the Widom-601 sequence corresponds to the ***R_g_*** of DNA around 45 Å with further unwrapping, a neighboring minimum appears at 49 Å and 51 Å (**Figure 2B**). Similar to the ASP sequence, the Widom-601 sequence appears to be unwrapping at the left side of the NCP (601-L, TA-rich side), consistent with prior MD simulations^56^ and nuclease digestion assays^49^. For H2A.Z variant, the first minimum in the PMF corresponds to the ***R_g_*** of DNA around 47 Å with further unwrapping, a neighboring minimum appears at 49 Å and 51 Å (**Figure 2C**). The histone H2A.Z variant system shows right-side unwrapping of the nucleosome (601-R, TA-poor side) as compared to the ASP and Widom-601 systems. To assess the robustness of these results with respect to salt concentration, we performed additional atomistic umbrella sampling simulations at 2.4 M NaCl for the ASP and Widom-601 systems, matching the ionic conditions of our earlier unbiased Anton simulations^56^. The high-salt PMF profiles preserve the same directional preferences and qualitative free energy features observed at physiological salt, confirming that the unwrapping asymmetry is an intrinsic property of each sequence rather than a consequence of ionic screening (**Figure S1**).

**Figure 2.**
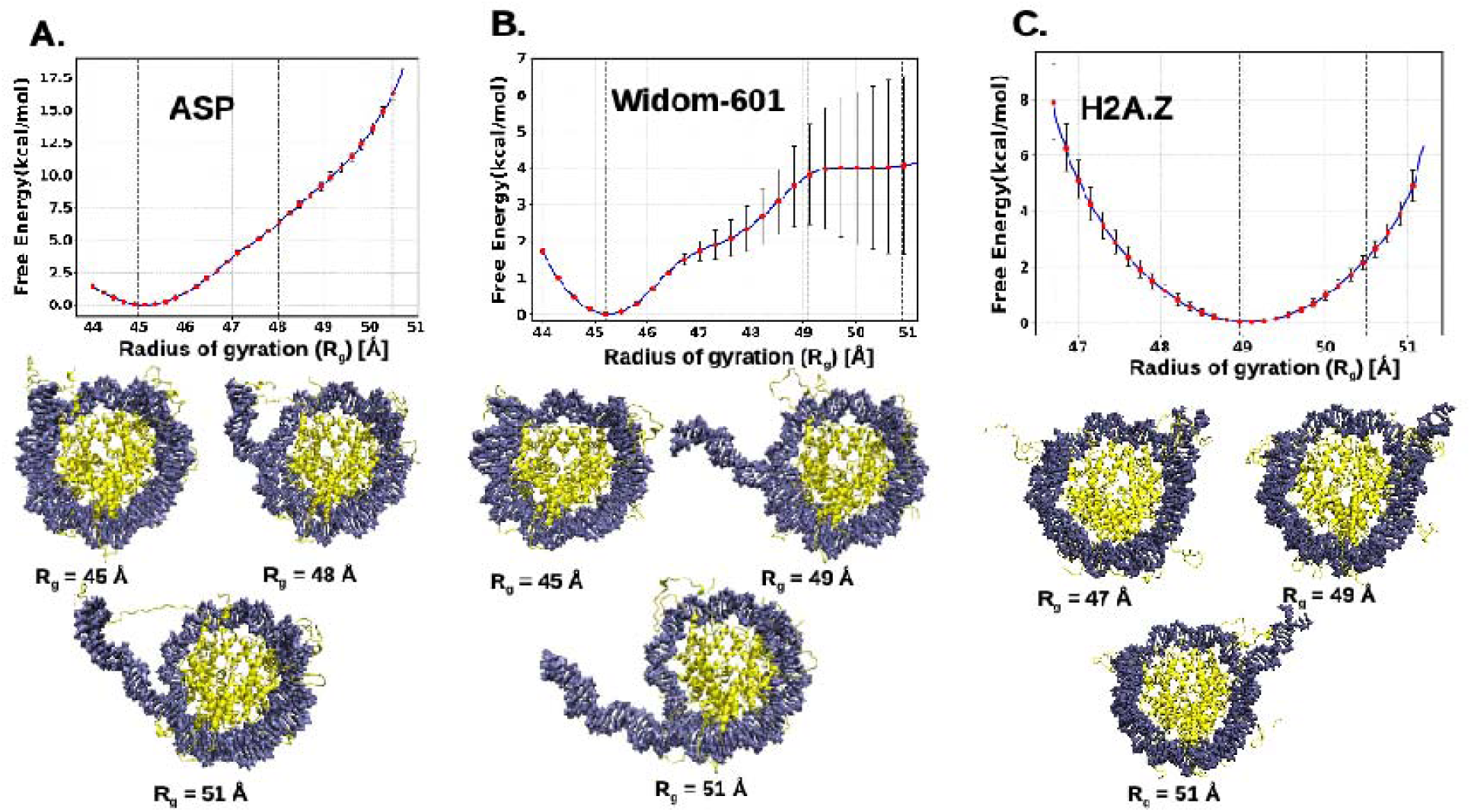
Free energy profile as a function of Radius of Gyration (R_g_). **(a)** The potential mean force (PMF) plot displays a blue curve representing the average free energy profile with error bars for the ASP sequence. The vertical dotted lines indicate the selected conformations shown below the PMF plot, at a radius of gyration of 45 Å, 48 Å, and 51 Å. **(b)** The PMF plot displays a blue curve representing the average free energy profile with error bars for the Widom-601 sequence. The vertical dotted lines indicate the selected conformations shown below the PMF plot, at a radius of gyration of 45 Å, 49 Å, and 51 Å. **(C)** The PMF plot shows a blue curve representing the average free-energy profile, with error bars, for the H2A.Z variant. The vertical dotted lines indicate the selected conformations shown below the PMF plot, at a radius of gyration of 47 Å, 49 Å, and 51 Å.

### 3.2. Histone H3 N-terminal tail modulates the DNA unwrapping

In the first section, we showed that DNA unwrapping pathways are sequence dependent **(Figure 2A–B)** and influenced by histone variant **(Figure 2B–C)**. It has been reported earlier that H3 N-terminal tails are important to facilitate breathing motion (less pronounced DNA unwrapping typically over 5-10bp) in the canonical nucleosome^56^. Here, we calculate the number of contacts (N_c_) between DNA ends (End1 and End2) with the N-terminal H3 tails. **Figure 3A** shows the position of both H3 histone monomers along with the tails. DNA ends are denoted as End1, marked in red, and End2, marked in blue. Both ends are approximately 30 bp in length. Both End1 and End2 are interacting with N-terminal H3 histone tails, Tail1 and Tail2, respectively. We calculate the average number of contacts in each window obtained from US simulations as discussed earlier so that we can identify the change of average contacts as a function of *R_g_*. For the ASP, we show that the N-terminal H3 tails are in contact with DNA in each window (see **Figure 3B**), although the number of contacts differs between the two ends. In most windows, Tail1 makes more contacts with DNA End1 (red) compared to the other end. The contact-based analysis shows different behaviors for the Widom-601 system. We find that the N-terminal H3 Tail1 interacts with DNA End1 throughout the windows, whereas the contact between DNA End2 and the N-terminal H3 Tail 2 decreases as *R_g_* increases. After *R_g_* = 48 Å, there are no contacts between DNA and N-terminal H3 Tail2 (see **Figure 3C**). The contacts between the tail and the DNA ends remains present across all windows in the case of the histone variant H2A.Z (see **Figure 3D**). However, the average number of contacts is generally higher for the interaction between H3 Tail1 and DNA End1 (marked in red, **Figure 3D**).

**Figure 3.**
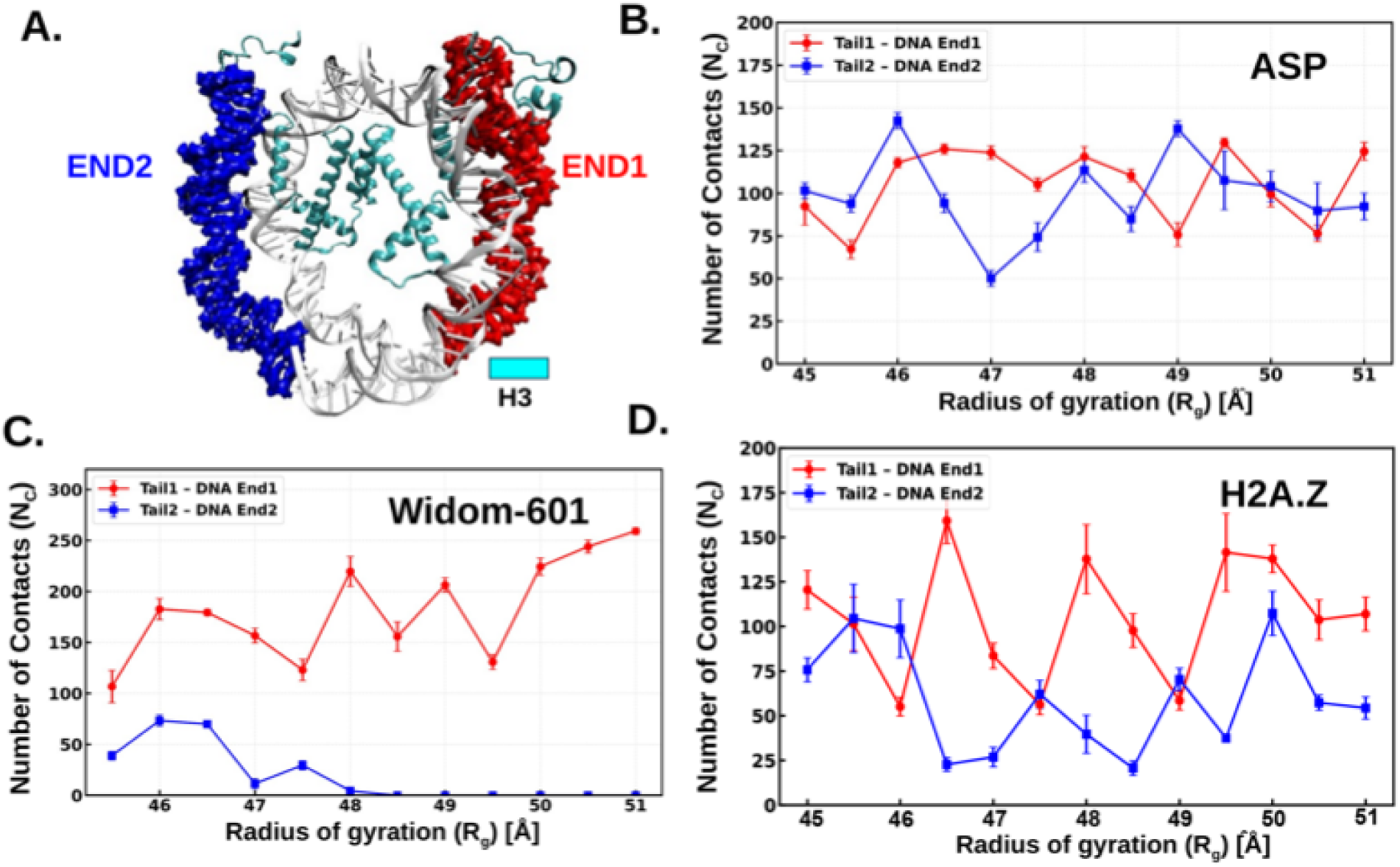
The number of contacts (Nc) between histone H3 N-terminal and DNA ends. **(A)** The NCP structure shows histone H3 (cyan) and two DNA ends: DNA end1 in red and DNA end2 in blue, both approximately 30 base pairs in length. **(B)** The average number of contacts between the H3 N-terminal tail and the DNA ends is shown as a function of the radius of gyration for the ASP sequence across each umbrella sampling window. **(C)** The average number of contacts between the H3 N-terminal tail and the DNA ends is shown as a function of the radius of gyration for the Widom-601 sequence across each umbrella sampling window. **(D)** The average number of contacts between the H3 N-terminal tail and the DNA ends is shown as a function of the radius of gyration for the histone H2A.Z variant across each umbrella sampling window. The distance cut-off for contacts is 4.5 Å for all the systems.

### 3.3. Structural characterization of DNA unwrapping

To further quantify the extent of DNA unwrapping suggested by the tail-DNA contact analysis in the previous section, we next examine the distance between DNA ends and the dyad (SHL0, see **Figure 1A**). The increase in this distance quantifies progressive unwrapping from the entry–exit regions toward the dyad. We calculate the distance in each window and plot as a function of different *R_g_* values. **Figure 4A** shows the distances for DNA ends in the ASP system, where SHL0-DNA End1 is represented in red, while the other end is depicted in blue. DNA End2 shows an increase in distance from 37 Å to 75 Å, whereas End1 does not change significantly across different *R_g_* values, remaining around 37 Å. This indicates asymmetric unwrapping of the nucleosome ends, as reported in several previous studies^96^. The Widom-601 sequence similarly shows asymmetric unwrapping; however, the distance between DNA End2 and SHL0 is significantly higher compared to the ASP sequence, reaching up to around 110 Å (**Figure 4B**).

**Figure 4.**
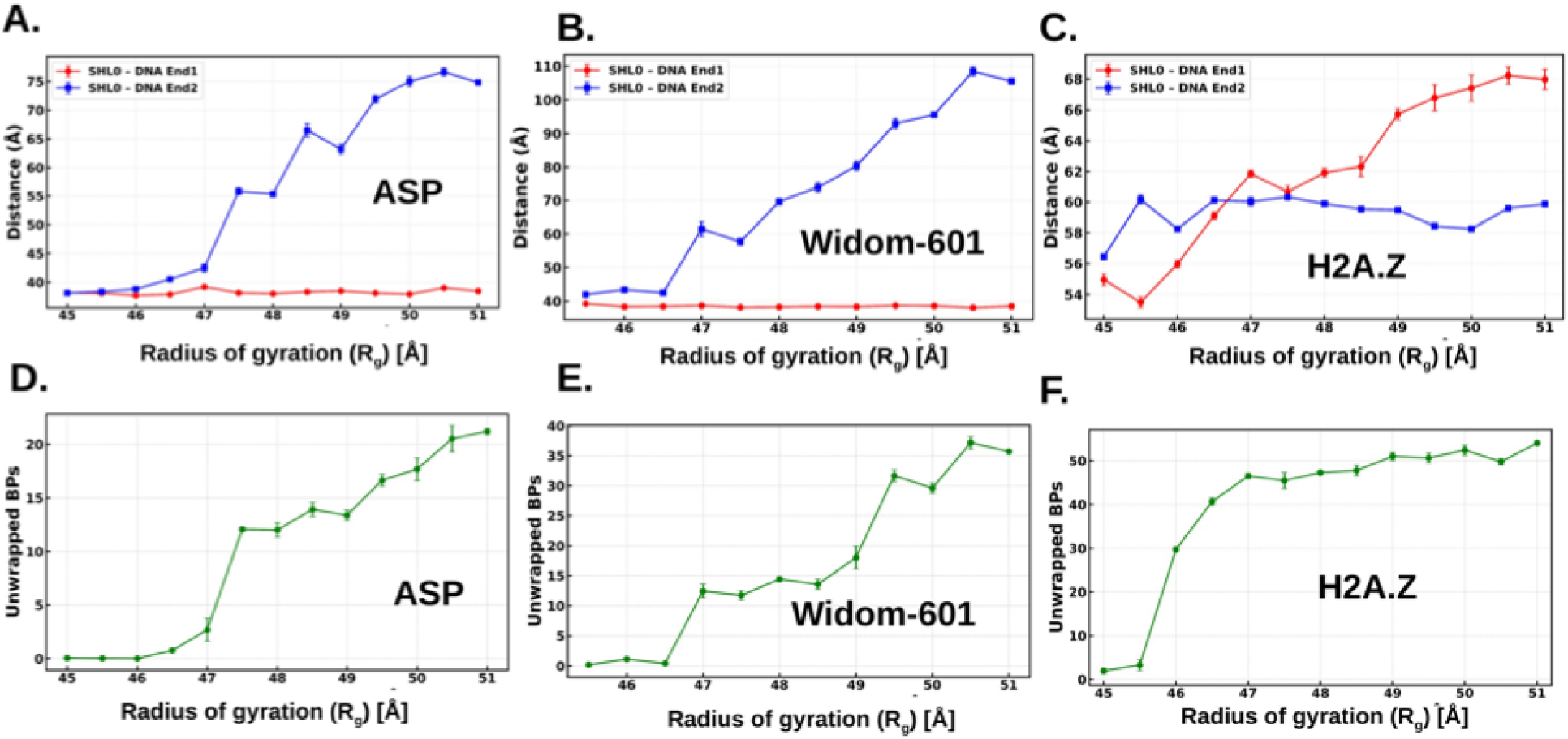
The distance between SHL0 and DNA ends, as well as the unwrapping of DNA base pairs. The distance between the SHL0 and DNA ends is calculated for each umbrella sampling window. **(A)** The average distance between SHL0 and DNA end 1 (red) as well as DNA end 2 (blue) for the ASP sequence as a function of radius of gyration for each umbrella sampling window is shown, indicating the increase in the distance for DNA end 1. **(B)** The average distance between SHL0 and DNA end 1 (red) as well as DNA end 2 (blue) for the Widom-601 sequence as a function of radius of gyration for each umbrella sampling window is shown, indicating the increase in the distance for DNA end 1. **(C)** The average distance between SHL0 and DNA end 1 (red) as well as DNA end 2 (blue) for the H2A.Z variant as a function of radius of gyration for each umbrella sampling window is shown, indicating the increase in the distance for DNA end 2. **(D)** The DNA unwrapping base pairs as a function of radius of gyration shows increase in the unwrapping across the radius of gyration reaction coordinate for ASP sequence, **(E)** for Widom-601 sequence and **(F)** for histone H2A.Z variant.

Interestingly, the histone variant system H2A.Z shows the opposite behavior, where DNA End1 exhibits a larger breathing distance compared to the other end as a function of *R_g_* (**Figure 4C**). Overall, our simulation study indicates that changing the DNA sequence modulates the extent and asymmetry of nucleosome unwrapping. In contrast, altering the histone variant can reverse the preferred direction of unwrapping between the two DNA ends. Together, these results highlight a coupled dependence of DNA sequence and histone variant on nucleosome unwrapping dynamics.

We further quantify DNA unwrapping in terms of the number of unwrapped base pairs (bps) from both ends. **Figure 4D** shows the number of unwrapped bps for the ASP system at different *R_g_*values. Up to *R_g_* = 47 Å, the number of unwrapped bps remains around 5; however, beyond this point, it shows a sharp increase to around 13. As *R_g_* increases further, the total number of unwrapped bps reaches approximately 20. For the Widom-601 sequence, the trend is different. Up to *R_g_* = 46.5 Å, the number of unwrapped bps does not change significantly, but beyond this point, there is a sharp increase (see **Figure 4E**). It then remains relatively plateaued with further increases in *R_g_*, eventually reaching around 35 bps at *R_g_* = 51 Å. The histone variant system H2A.Z shows yet another distinct behavior (see **Figure 4F**). The number of unwrapped bps increases sharply at *R_g_* = 47 Å, after which it does not change significantly with further increases in *R_g_* . Overall, this study indicates that DNA sequence and histone variants differentially regulate the extent and progression of nucleosome unwrapping.

### 3.4. Nucleosome unwrapping using the SIRAH coarse-grained force field

Next, we examine the ability of SIRAH coarse-grained force field to capture nucleosome unwrapping dynamics and assess how faithfully it reproduce the sequence and histone variant dependent unwrapping pathways observed in atomistic simulations.

**Figure 5A** shows the PMF profile for the ASP nucleosome obtained from umbrella sampling using the SIRAH CG force field. We identify representative structures from two regions of the PMF: one at *R_g_* = 46 Å, corresponding to the minimum, and another at *R_g_* = 47 Å, indicating DNA unwrapping from the left side (End2). For the Widom-601 system (**Figure 5B**), the PMF minimum is slightly shifted to *R_g_* = 46 Å, while a structure at *R_g_* = 51 Åshows DNA unwrapping from the same side as observed in the ASP system. For the H2A.Z variant, the PMF minimum is further shifted to *R_g_* = 48 Å(**Figure 5C**). A structure at *R_g_* = 50 Åindicates unwrapping occurring from both DNA ends, although the right end (End1) shows more pronounced unwrapping. Overall, the SIRAH CG simulations qualitatively reproduce the sequence- and histone variant–dependent unwrapping pathways observed in atomistic simulations, including the preferred direction of unwrapping. However, subtle differences in the PMF minima and the extent of unwrapping suggest limitations in capturing fine energetic details at the coarse-grained level.

**Figure 5.**
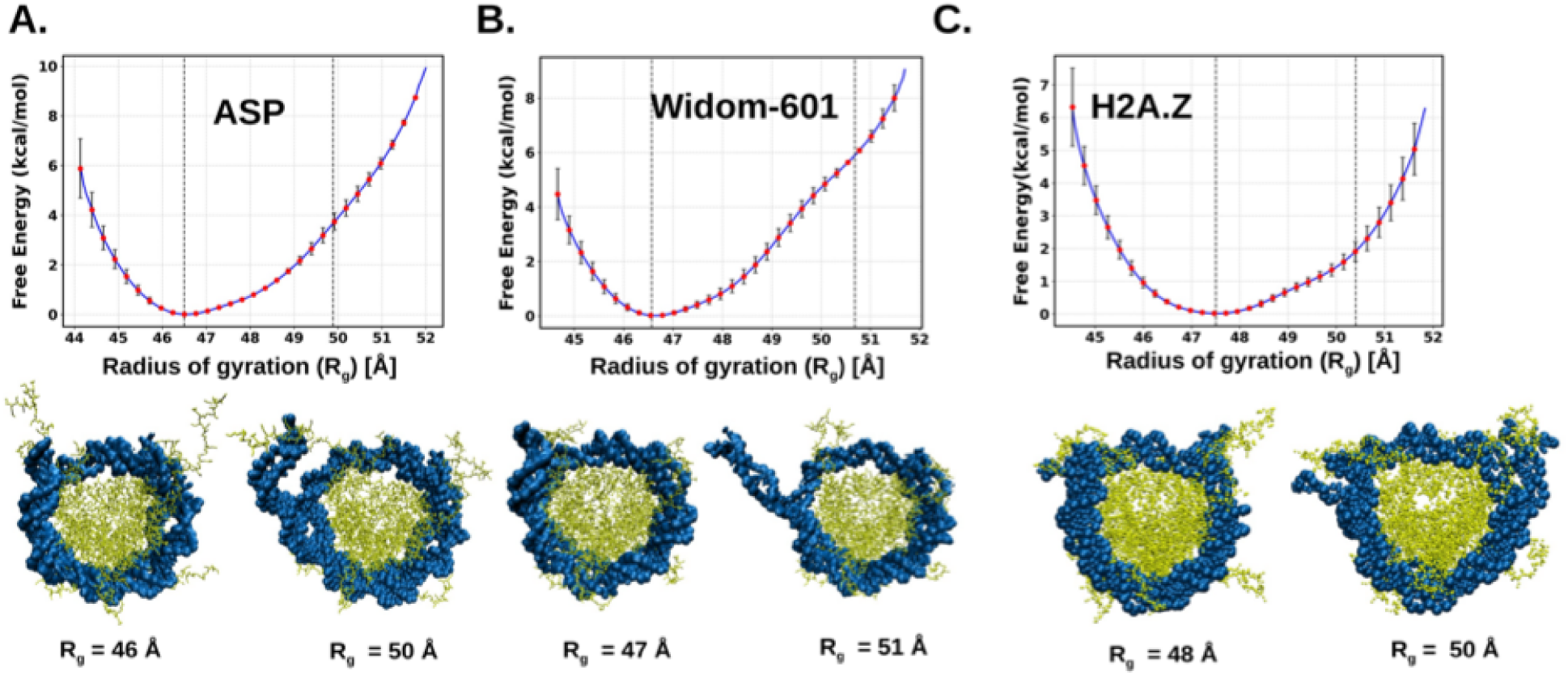
Free energy profile as a function of Radius of Gyration (R_g_) based on SIRAH Coarse-grained simulations. **(A)** The potential mean force (PMF) plot displays a blue curve representing the average free energy profile with error bars for the ASP sequence. The vertical dotted lines indicate the selected conformations shown below the PMF plot, at a radius of gyration of 47Å and 50 Å. **(B)** The PMF plot displays a blue curve representing the average free energy profile with error bars for the Widom-601 sequence. The vertical dotted lines indicate the selected conformations shown below the PMF plot, at a radius of gyration of 47 Å and 51Å. **(C)** The PMF plot shows a blue curve representing the average free-energy profile, with error bars, for the H2A.Z variant. The vertical dotted lines indicate the selected conformations shown below the PMF plot, at a radius of gyration of 48 Å and 51 Å.

### 3.5. Simulated H3 tail flexibility is consistent with NMR-detected tail dynamics

To assess whether the H3 N-terminal tail dynamics captured by our atomistic simulations are consistent with experiment, we compared the simulated per-residue change in flexibility of Tail1 upon H2A.Z incorporation (ΔRMSF = RMSF_H2A.Z_ − RMSF_H2A_) with published solution NMR amide peak intensity ratios (*I_H2A.Z_*/*I_H2A_*) reported for two Widom-601 nucleosomes bearing Sox2-Oct4 composite motifs derived from the mouse Fgf4 (601-62_F_) and Nanog (601-62_N_) promoter regions (**Figure 6A**).^49^ Because amide intensity reflects the net transverse relaxation, it is determined by two competing contributions: faster picosecond-nanosecond (*ps-ns*) motion narrows resonances and raises intensity, whereas microsecond-millisecond (*µs-ms*) conformational exchange broadens them and lowers it. Since ΔRMSF is computed over the nanosecond-scale simulation window, it captures only the fast (*ps-ns*) amplitude of motion and is insensitive to *µs-ms* exchange. Therefore, a residue that becomes more mobile upon H2A.Z incorporation is expected to show both a larger ΔRMSF and a higher intensity ratio, except where a slower exchange process dominates its relaxation and instead attenuates the resonance or where the fast-narrowing response saturates and the intensity becomes insensitive to further changes in flexibility (**Figure 6A**).

**Figure 6.**
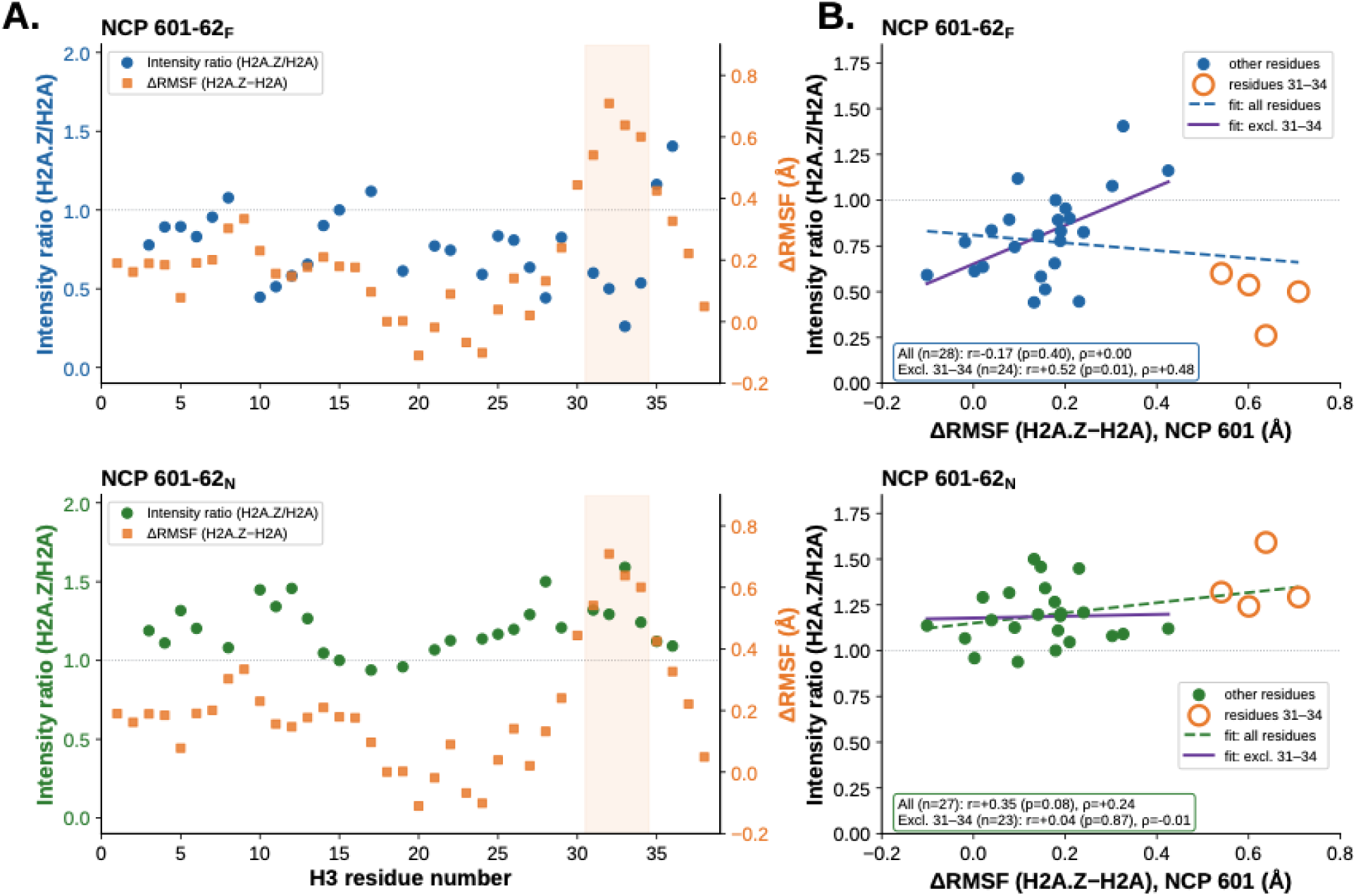
Comparison of the simulated ΔRMSF with published NMR amide intensity ratios for the H3 N-terminal tail. **(A)** Per-residue H2A.Z/H2A amide intensity ratio (left axis) and simulated ΔRMSF (H2A.Z − H2A; orange squares, right axis) for the 601-62_F_ (top) and 601-62_N_ (bottom) constructs; the shaded band marks residues 31-34. **(B)** Intensity ratio versus simulated ΔRMSF for 601-62_F_ (left) and 601-62_N_ (right); open orange symbols denote residues 31-34, dashed lines are linear fits to all resolved residues and solid lines are fits excluding residues 31-34, with Pearson (*r*) and Spearman (ρ) coefficients annotated. In 601-62_F_, residues 31-34 invert an otherwise positive relationship, whereas in 601-62_N_ they follow the fast-motion trend and account for the entire weak positive correlation.

For the 601-62_F_ construct, ΔRMSF and the intensity ratio were uncorrelated across the full tail (Pearson *r* = −0.17, *p* = 0.40; **Figure 6A-B**). Excluding residues 31-34, however, a significant positive correlation emerged (*r* = +0.52, *p* = 0.009; Spearman ρ = +0.48, *p* = 0.019), as anticipated for faster *ps-ns* motional behavior (**Figure 6B**). Residues 31-34 departed from this trend, exhibiting both the largest ΔRMSF values (up to ∼0.7 Å) and the lowest intensity ratios (mean ≈ 0.48). Since increased fast-timescale mobility would sharpen rather than weaken these resonances, the suppressed intensity at residues 31-34 implicates a *µs-ms* exchange process that dominates their transverse relaxation. Prior NMR relaxation studies^39, 65^ reporting on *ps-ns* motions have characterized these residues (A31-T32-G33-G34) as a flexible hinge N-terminal to the basic DNA-anchoring segment (K36/K37, R40/R42) mapped by MD simulations^97^, a pivotal position that would make them especially sensitive to a slow conformational switch of the adjacent anchor. For the 601-62_N_ construct, the intensity ratios were uniformly at or above unity across the entire tail (∼0.9-1.6), including residues 31-34 (mean ≈ 1.36) (**Figure 6A**). The weak overall correlation (r = +0.35, p = 0.08) with ΔRMSF was carried almost entirely by these residues and abolished upon their exclusion (r = +0.04, p = 0.87) (**Figure 6B**), consistent with intensities near the motional-narrowing limit. In contrast to the systematically lower 601-62_F_ ratios (mostly <1), these uniformly high intensities indicate that *µs-ms* exchange likely broadens the entire 601-62_F_ tail in the presence of H2A.Z while contributing little in 601-62_N_, where fast-timescale narrowing governs relaxation.

Because both constructs share the same simulated ΔRMSF profile, the uniformly lower 601-62_F_ intensities, most pronounced at residues 31-34, reflect sequence-dependent slow dynamics induced by H2A.Z rather than differences in fast-timescale dynamics, while the positive ΔRMSF-intensity correlation elsewhere in 601-62_F_ corroborates the simulated fast flexibility of the tail. Since the two constructs differ only in the Sox2-Oct4 composite motif at the nucleosome edge where the H3 tail contacts DNA, the local sequence appears to encode both the fast flexibility and slower DNA engagement of the tail, offering a route for sequence and H2A.Z to jointly tune nucleosome accessibility at pioneer factor sites.

## 4. Discussion

The results presented here contribute to a growing understanding of how nucleosome composition encodes chromatin accessibility. A recent large-scale coarse-grained study by *Collepardo-Guevara et al.*^98^ examined force-induced unwrapping across 40 nucleosomes spanning genomic sequences, histone variants, and post-translational modifications, finding that unwrapping barriers arise from topologically protected partially unwrapped intermediates governed by DNA flexibility and histone–DNA electrostatics, with histone variant and modification effects operating in a strongly non-additive manner. The present study is complementary in scope: where that work surveys breadth across many sequences and modifications, we probe the atomistic mechanism for three specific nucleosome systems and directly validate a CG force field against those benchmarks. Our finding that progressive H3 tail–DNA disengagement acts as a directional switch governing which end unwraps preferentially provides a residue-level mechanistic basis that complements the electrostatic hierarchy identified by Collepardo and colleagues. Furthermore, the reversal of unwrapping directionality upon H2A.Z substitution points to context-dependent effects of histone variant substitution on unwrapping pathway geometry, not just energetics, underscoring the value of atomistic resolution for mechanistic interpretation.

Earlier microsecond-long unbiased atomistic simulations^56^ and coarse-grained studies^57^ for these sequences demonstrated sequence-dependent nucleosome dynamics, capturing breathing motions. Breathing motion was more pronounced in the Widom-601 sequence than in ASP in atomistic simulations and was initiated preferentially from the 601L end, which corresponds to End2 in the present study. Consistently, our SMD simulations reveal preferential unwrapping from End2 for the Widom-601 sequence, corroborating this earlier observation. The SIRAH coarse-grained SMD simulations establish the same preferential unwrapping from End2, consistent with the preferential breathing observed at End2 in earlier unbiased SIRAH CG simulations of the Widom-601 system.

Interestingly, while breathing motion was not prominent for the ASP sequence in atomistic unbiased simulations, earlier unbiased SIRAH coarse-grained simulations captured asymmetric breathing motion, with End2 exhibiting greater flexibility than End1. Notably, the present SMD and umbrella sampling simulations reveal that unwrapping also initiates preferentially from the same End2 in the ASP system. The consistency of the preferred DNA end across both breathing and unwrapping regimes and across two distinct simulation approaches suggests that this directional preference is an intrinsic sequence-encoded property of each nucleosome system, and that local breathing motion may serve as a structural precursor to large-scale unwrapping. In the present study, consistent asymmetric unwrapping is observed for both DNA sequences, as evidenced by markedly greater SHL0-End2 distances compared to End1 across umbrella sampling windows, providing a unified geometric basis for sequence-dependent unwrapping asymmetry at both atomistic and coarse-grained levels.

Based on previous all-atom simulation studies and combined NMR-MD experiments, the H2A.Z variant promotes substantially greater DNA unwrapping and nucleosome breathing than the canonical H2A nucleosome, increasing DNA accessibility. The H2A.Z C-terminal tail is shorter and less positively charged, leading to fewer and more transient contacts with DNA at the nucleosomal entry/exit regions.^49, 99^ Importantly, recent work combining NMR spectroscopy and MD simulations demonstrated that H2A.Z incorporation disrupts an H3 R42–H2A E121 salt bridge, leading to more disordered H3 N-terminal tail conformations with fewer contacts bridging the two DNA gyres, particularly at SHL±2.^49^ This H3 tail disengagement was shown to directly increase DNA accessibility and facilitate pioneer factor binding at both end-positioned and internal nucleosome sites.^49^ The present study extends these findings into the context of enhanced sampling free energy calculations, showing that the shift in preferred unwrapping direction with H2A.Z is mechanistically consistent with altered H3 tail–DNA contacts observed in unbiased simulations. Independent NMR measurements of site-specific electrostatic potentials across all four core histone tails in the intact nucleosome core particle provide additional, orthogonal support for treating tail-DNA engagement as an electrostatically governed switch^100, 101^.

These analyses describe the altered H3 tail-DNA engagement structurally but do not resolve the timescales on which the tail moves. To add this dimension, we compared our simulated ΔRMSF with published NMR intensities^49^, which provides an experimental cross-check on the H3 tail dynamics and points to an additional, slower motional component. Because both amide intensity and ΔRMSF report on the fast *ps-ns* motions, their agreement outside residues 31-34 in 601-62_F_ indicates that the simulations capture the fast-timescale response of the tail to H2A.Z. Amide intensity, however, additionally reports *µs-ms* conformational exchange, to which ΔRMSF is insensitive. Residues at which the two measurements diverge, therefore, flag positions where such slow motions may contribute. On this basis, we tentatively attribute the large intensity attenuation at residues 31-34 in 601-62_F_ to a sequence-dependent *µs-ms* process, possibly an interconversion of the tail between DNA-bound and released states, that is present in the Fgf4-derived sequence but diminished, or too fast to broaden the resonances, in the Nanog-derived 601-62_N_ sequence. We emphasize that this assignment is speculative, being inferred from the opposing behavior of the two observables rather than measured directly. Its confirmation will require NMR experiments that report explicitly on *µs-ms* dynamics, such as relaxation dispersion (CPMG and *R*1ρ) or CEST measurements,^102^ together with controls for amide hydrogen exchange.^103^ More broadly, our simulated ΔRMSF and NMR intensity agree in the fast-motion regime but diverge where slow exchange is implicated, offering a practical way to corroborate the simulated fast-timescale flexibility and identify candidate slow-motion residues of the H3 tail for targeted experimental study. A promising complement to the RMSF-based proxy used here would be to directly back-calculate NMR relaxation observables (R1, R2, and rotational correlation times) from the simulated trajectories using frameworks such as MD2NMR^104^, enabling a more direct, quantitative comparison to experimental relaxation data than the RMSF-intensity correlation used in this study.

The H3 N-terminal tails play a critical role in modulating nucleosome stability and DNA dynamics. Earlier experimental studies^105, 106^ have shown that removal of H3 tails increases DNA accessibility and unwrapping rates, highlighting their importance in maintaining nucleosome integrity. Computational studies^60, 107^ have similarly demonstrated that H3 tail-DNA contacts are dynamically coupled to breathing motions and unwrapping events. Our earlier microsecond-timescale unbiased atomistic simulations of the same Widom-601 sequence^56^ identified a three-step mechanism underlying this coupling: the H3 N-terminal tail first condenses on the minor groove near SHL±6, then extends to condense on SHL±7, and finally forms an α-helix that docks onto the H3 core α-N helix, pulling the tail, and the associated DNA entry/exit region, away from the histone surface; this helix-helix linkage step occurred preferentially at the more mobile DNA end, consistent with the asymmetric H3 tail-DNA disengagement reported below. Independent solution NMR studies of H3 tail dynamics in reconstituted nucleosomes and nucleosome arrays similarly show that H3 tail-DNA contact strength is a graded, modulatable property rather than a fixed on/off state, directly paralleling the progressive disengagement described here^37–39^. Consistent with this framework, a recent combined NMR and MD study of the H3.3 G34R cancer mutation showed that a single substitution in the H3 tail hinge region shifts its conformational ensemble toward increased DNA association and decreased local dynamics, the converse of the H2A.Z-induced disengagement described below, further supporting H3 tail-DNA engagement as a bidirectionally tunable, disease-relevant switch.^108^ In the present study, we observe a strong sequence-dependent tail-DNA interaction during DNA unwrapping from the nucleosome (see **Figure 3**). For the ASP system, both DNA ends maintain balanced interactions with their respective H3 tails throughout the unwrapping process. However, despite this balanced tail engagement, End2 exhibits more pronounced unwrapping during SMD simulations compared to End1, suggesting that tail-DNA contacts alone do not fully determine the directionality of unwrapping in this sequence. This is consistent with our earlier unbiased simulations of the ASP system^56^, which showed that ASP nucleosome dynamics is instead governed by a distinct “wave-like” mechanism involving cooperative condensation of the H2A C-terminal and H2B N-terminal tails at SHL±7 and SHL±5, respectively, rather than by H3 N-terminal tail engagement, explaining why H3 tail-DNA contacts alone do not predict unwrapping directionality for this sequence. In contrast, for the Widom-601 nucleosome, End2 shows progressive disengagement from H3 Tail2 beyond R_g_ = 48Å, providing a direct molecular basis for its preferential unwrapping. This finding is consistent with single-molecule FRET studies demonstrating asymmetric unwrapping of the Widom-601 nucleosome and with the observation by *Ngo et al.*^29^ that one DNA end interacts more strongly with the histone core under tension. Our contact analysis thus provides a microscopic explanation for this experimentally observed asymmetry, identifying H3 tail disengagement as the structural basis of directional unwrapping preference in the Widom-601 nucleosome.

The PMF profiles obtained from atomistic umbrella sampling simulations reveal distinct free energy landscapes across the three nucleosome systems. The ASP and H2A.Z nucleosomes each exhibit a single well-defined minimum, reflecting a relatively smooth unwrapping pathway, whereas the Widom-601 nucleosome displays an additional local minimum, indicative of a metastable intermediate state along the unwrapping pathway. Notably, the total free energy cost of unwrapping is higher for the ASP system compared to Widom-601, suggesting that despite Widom-601 being a strongly positioning sequence, the ASP nucleosome presents a higher energetic barrier to complete unwrapping. This is consistent with the base pair-level elastic deformation analysis from our earlier unbiased simulations of these sequences^56^, which found lower dinucleotide force constants, indicating greater local deformability, in the Widom-601 entry/exit regions (SHL±7) relative to the corresponding ASP regions, providing an independent physical basis for the comparatively lower free energy cost of unwrapping in Widom-601. Partially destabilized nucleosome states consistent with such metastable intermediates have also been detected in solution by methyl-TROSY NMR, providing experimental corroboration that intermediate states are not simulation artifacts^101^. The SIRAH coarse-grained simulations qualitatively reproduce these features including the higher free energy cost for ASP relative to Widom-601 confirming the ability of the SIRAH force field to capture sequence-dependent free energy landscapes. However, quantitative differences in absolute barrier heights between atomistic and SIRAH simulations persist across all systems, reflecting the inherent limitations of coarse-grained representations in accurately capturing the energetics of protein-DNA interactions. These discrepancies likely arise from several compounding factors: the reduced degrees of freedom inherent to the SIRAH mapping may underrepresent the configurational entropy of the disordered histone tails, since a coarser resolution can restrict the range of tail conformations sampled; the simplified treatment of electrostatics and solvation in the coarse-grained ion and water models may not fully capture the fine electrostatic balance underlying specific contacts such as the H3 R42-H2A E121 salt bridge implicated in H2A.Z-dependent unwrapping; and the current SIRAH parameters were developed and validated primarily against structural benchmarks rather than protein-DNA unwrapping free energy landscapes directly. Reparameterizing SIRAH nonbonded interactions for histone tail-DNA contacts against atomistic free energy references of the kind reported here, together with systematic back-mapping and reweighting validation, represents a promising route to improving the quantitative accuracy of the force field for future large-scale chromatin simulations. For the H2A.Z system, the quantitative agreement between atomistic and SIRAH PMF profiles is notably closer compared to the canonical systems, suggesting that the altered histone-DNA interface in H2A.Z nucleosomes may be more faithfully represented at the coarse-grained level. Together, these results establish SIRAH as a reliable tool for qualitative exploration of sequence- and variant-dependent nucleosome unwrapping landscapes, while highlighting the continued necessity of atomistic simulations for precise free energy quantification.

## 5. Conclusions

In summary, we have performed a comprehensive multiscale simulation study of nucleosome unwrapping by combining atomistic and SIRAH coarse-grained umbrella sampling simulations across three nucleosome systems. Our results demonstrate that nucleosome unwrapping is inherently asymmetric and strongly sequence-dependent, with distinct free energy landscapes observed for ASP and Widom-601 nucleosome. Tail-DNA contact reveals that progressive tail disengagement from the preferred DNA end acts as a molecular determinant for DNA unwrapping which establish a mechanistic link between tail-DNA interactions and unwrapping directionality. The variation in histone, H2A.Z shows modified DNA-tail interaction by altering the preferred unwrapping direction. The observations based on atomistic simulations are further corroborated by SIRAH coarse-grained simulations, which successfully reproduce the qualitative features of sequence- and variant-dependent unwrapping pathways, including the preferred unwrapping end and asymmetric behaviour. Although quantitative differences in free energy barrier heights persist in SIRAH simulations, highlighting the need for further refinement of coarse-grained interaction parameters to accurately capture the underlying energetics, the present study establishes SIRAH as a computationally efficient and reliable framework for investigating nucleosome unwrapping making it a promising tool for large-scale chromatin simulations, including nucleosomal arrays and longer timescale unwrapping events, that remain computationally demanding at the atomistic level. Finally, comparison of the simulated H3 N-terminal tail flexibility with published solution-NMR amide intensities corroborated the fast-timescale tail dynamics and pointed to a sequence-dependent *µs–ms* process, most pronounced at the tail-core junction (residues 31–34), underscoring how the local DNA sequence tunes both the fast flexibility and slower DNA-engagement of the H3 tail. Together, these findings provide molecular insight into how DNA sequence and histone variant composition jointly govern nucleosome unwrapping dynamics, with important implications for understanding chromatin accessibility and gene regulation.

## Supporting Information

Summary table for system setup (Table S1). US results for higher salt concentration in Figure S1.

## Acknowledgments

This work was supported by grants from the NIH, 1R15GM146228-01 awarded to S.M.L. and R01GM147642 awarded to E.N.N. R. P. and T. R. are grateful for support from The Rosemary O’Halloran Scholarship. T. R. was also supported through an NSF NRT grant DGE-2151945. This work used Bridges-2 at the Pittsburgh Supercomputing Center through allocation BIO230066 from the Advanced Cyberinfrastructure Coordination Ecosystem: Services & Support (ACCESS) program, which is supported by NSF grants 2138259, 2138286, 2138307, 2137603, and 2138296. We thank Dr. Ananya Majumdar, director of the Biomolecular NMR center (JHU), for assistance with NMR data collection. We are grateful for discussions with Prof. Sergio Pantano.

## Data Availability

Analysis codes will be made available on a GitHub Repository.

## Notes

### Competing Interest Statement

The authors have declared no competing interest.

### Summary of Updates

Section including corroborating NMR experiments is added to the manuscript, as well as additional co-author

