## Supplementary Information for "DNA Sequence and Histone Variant H2A.Z Jointly Govern Nucleosome Unwrapping Pathways"

^5^Currently at: Institute of Nanotechnology, Karlsruhe Institute of Technology (KIT), [Kaiserstraße 12, 76131 Karlsruhe](https://www.google.com/maps/place/data=!4m2!3m1!1s0x4797064d7da356a3:0xbdaeb296b0747bed?sa=X&ved=1t:8290&ictx=111)

**Supporting Information:**

Table S1. Summary of initial set-up of the atomistic simulations

| **NCP systems** | **ASP (AA)** | **Widom601 (AA)** | **H2A.Z (AA)** | **ASP (CG)** | **Widom601 (CG)** | **H2A.Z (CG)** |
| --- | --- | --- | --- | --- | --- | --- |
| Box dimensions, Å | 159 x 191 x 112 | 171 x185 x 124 | 171 x 127 x 185 | 212 x 212 x 212 | 213 x 213 x 213 | 171 x 127 x 185 |
| No. of atoms | 444888 | 448776 | 459742 | 114459 | 117837 | 459742 |
| No. of solvent | 104740 | 105816 | 108512 | 26587 | 27448 | 108512 |
| No. of Na^+^ ions | 472 | 486 | 496 | 896 | 930 | 496 |
| No. of Cl^-^ ions | 356 | 358 | 366 | 783 | 805 | 366 |
| No. of Mg^2+^ ions | 14 | 8 | 8 | 14 | 8 | 8 |
| Salt Conc. (M) | 0.15 | 0.15 | 0.15 | 0.15 | 0.15 | 0.15 |
| Change in Rg during SMD | 4.50 nm to 5.10 nm | 4.50 nm to 5.10 nm | 4.50 nm to 5.10 nm | 4.50 nm to 5.10 nm | 4.50 nm to 5.10 nm | 4.50 nm to 5.10 nm |
| Force constant,  k_r_ | 837 kJ mol^-1^ nm^-2^ | 837 kJ mol^-1^ nm^-2^ | 837 kJ mol^-1^ nm^-2^ | 837 kJ mol^-1^ nm^-2^ | 837 kJ mol^-1^ nm^-2^ | 837 kJ mol^-1^ nm^-2^ |
| Number of windows | 13 | 12 | 13 | 13 | 12 | 13 |

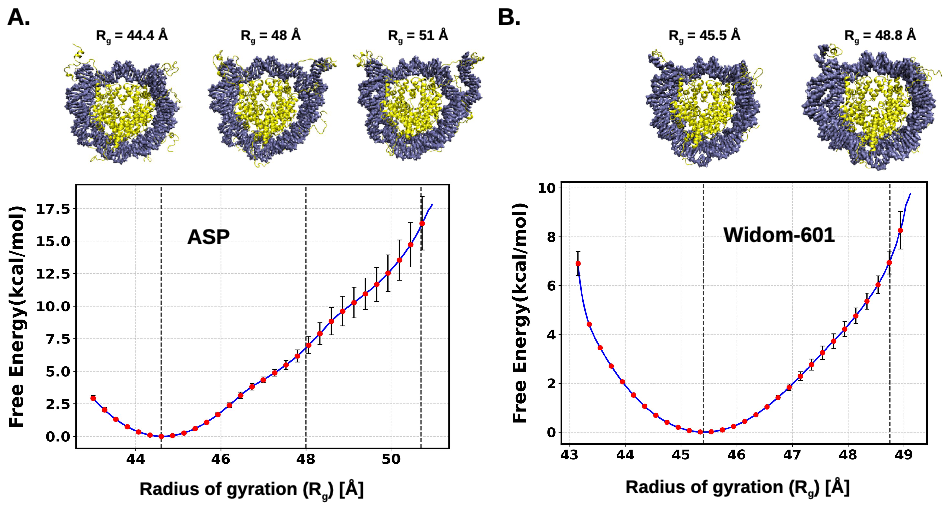
**Figure S1. Free energy profile as a function of Radius of Gyration (R_g_) in high salt (2.4 M) concentration using atomistic free energy calculation. (A)** The potential mean force (PMF) plot displays a blue curve representing the average free energy profile with error bars for the ASP sequence. The vertical dotted lines indicate the selected conformations shown below the PMF plot, at a radius of gyration of 44.4 Å, 48 Å, and 51 Å. **(B)** The PMF plot displays a blue curve representing the average free energy profile with error bars for the Widom-601 sequence. The vertical dotted lines indicate the selected conformations shown below the PMF plot, at a radius of gyration of 45.5 Å, 48.8 Å.
